# Whole genome bisulfite sequencing of Down syndrome brain reveals regional DNA hypermethylation and novel disorder insights

**DOI:** 10.1101/428482

**Authors:** Benjamin I. Laufer, Hyeyeon Hwang, Annie Vogel Ciernia, Charles E. Mordaunt, Janine M. LaSalle

## Abstract

Down Syndrome (DS) is the most common genetic cause of intellectual disability, in which an extra copy of human chromosome 21 (HSA21) affects regional DNA methylation profiles across the genome. Although DNA methylation has been previously examined at select regulatory regions across the genome in a variety of DS tissues and cells, differentially methylated regions (DMRs) have yet to be examined in an unbiased sequencing-based approach. Here, we present the first analysis of DMRs from whole genome bisulfite sequencing (WGBS) data of human DS and matched control brain, specifically frontal cortex. While no global differences in DNA methylation were observed, we identified 3,152 DS-DMRs across the entire genome, the majority of which were hypermethylated in DS. DS-DMRs were significantly enriched at CpG islands and de-enriched at specific gene body and regulatory regions. Functionally, the hypermethylated DS-DMRs were enriched for one-carbon metabolism, membrane transport, and glutamatergic synaptic signaling, while the hypomethylated DMRs were enriched for proline isomerization, glial immune response, and apoptosis. Furthermore, in a cross-tissue comparison to previous studies of DNA methylation from diverse DS tissues and reference epigenomes, hypermethylated DS-DMRs showed a strong cross-tissue concordance, while a more tissue-specific pattern was observed for the hypomethylated DS-DMRs. Overall, this approach highlights that low-coverage WGBS of clinical samples can identify epigenetic alterations to known biological pathways, which are potentially relevant to therapeutic treatments and include metabolic pathways. These results also provide new insights into the genome-wide effects of genetic alterations on DNA methylation profiles indicative of altered neurodevelopment and brain function.

## Introduction

Down Syndrome (DS), which is caused by Trisomy 21, results from an extra copy of human chromosome 21 (HSA21). DS is the most common genetic cause of intellectual disability and also the most common chromosomal aneuploidy in live-births, affecting approximately 1 in 691 in the United States.^1^ Maternal age is a risk factor for DS with a noticeable increase occurring in mothers greater than 35 years old.^2^ The prevalence of live births with DS has increased by 31% from 1979 through 2003.^3^ Life expectancy for people with DS has increased rapidly from 10 years in 1960 to 47 years in 2007.^4^ Overall, individuals with DS represent a substantial and growing portion of the population, suggesting a need for further study and therapeutic development for DS.

DS is characterized by distinct phenotypic traits, which include intellectual disability, facial dysmorphisms, short stature, congenital heart defects, and immune system abnormalities.^5^ Notably, DS is also associated with an increased risk for childhood leukemia, early onset Alzheimer’s disease, and a reduced occurrence of solid tumors.^6^ An understanding of the molecular mechanisms downstream of the genetic underpinnings of DS will also enable future therapeutic interventions as well as an improved understanding of the associated traits and disabilities in the non-DS population.

Although DS is the result of the increased copy number of a single chromosome, the regulation of gene expression is affected at a genome-wide level in a wide variety of tissues.^7,8^ While there is an increase in global gene expression from HSA21 in DS patients, not all HSA21 genes are differentially expressed, and differential gene expression is observed genome-wide across all chromosomes.^9–11^ An examination of fibroblasts from monozygotic twins discordant for DS revealed that differential gene expression occurred in distinct chromosomal domains that correlated with late DNA replication domains and lamina-associated domains (LADs).^12^ While the positioning of the LADs was not altered, H3K4me3, a histone modification associated with active promoters, was altered within the DS-associated chromosomal domains. Further investigation into fetal skin fibroblasts from additional sets of monozygotic twins uncovered differential DNA CpG methylation outside of HSA21.^13^ These regions were predominantly hypermethylated in DS, mapping to genes involved in embryonic organ morphogenesis. Reprogramming of the DS fibroblasts into induced pluripotent stem cells (iPSCs) revealed that select DS DMRs were maintained in the pluripotent state and correlated with differential gene expression and increased expression of the DNA methyltransferases DNMT3B and DNMT3L. Thus, the genome-wide differences seen in DS tissues are correlated with epigenetic modifications that may be responsible, in part, for the establishment and/or maintenance of differential expression of genes outside of HSA21 in DS.

Further investigations into the epigenetic mechanisms of DS have primarily focused on DNA methylation, where genome-wide differences in CpG methylation have been observed in a variety of DS tissues.^8^ Given the tissue-specific nature of epigenetic modifications, the differences have primarily been examined in cells and tissues relevant to DS phenotypes. For example, differential methylation has been observed in immune cell types in blood^14,15^ and fetal liver mononuclear cells^16^ at genes involved in the development of these cell types. Genome-wide methylation differences have also been observed in buccal epithelial cells, where some of differences were observed in genes related to cell adhesion, protein phosphorylation, and neurodevelopment.^17^ Notably, like neurons, buccal epithelial cells are derived from the ectoderm and demonstrate a more similar methylation pattern to the brain than blood.^18^ Overall, the differential CpG methylation typically appears to be related to genes involved in early developmental processes that are relevant to the cell type assayed.

While the CpG sites with differential methylation in DS are primarily specific to the tissue and cell types examined, most aberrantly methylated CpG sites in DS are hypermethylated. For example, an examination of DS placenta revealed global hypermethylation across all autosomes.^19^ Placental gene expression from HSA21 was upregulated, but genes outside of HSA21 with promoter hypermethylation in DS were down-regulated. Interestingly, the DS-differential genes had functions related to the immune system and neurodevelopment. Global CpG hypermethylation (<1% difference) was also present in both fetal (mid-gestation cerebrum) and adult (cerebellar folial cortex and frontal cortex) DS brain tissues as well as sorted neurons (NeuN+ nuclei) and glia (NeuN− nuclei) from adult frontal cortex.^20^ Only sorted T-lymphocytes (CD3+) from adult DS peripheral blood failed to demonstrate global hypermethylation. While global hypermethylation was observed in fetal and adult brain tissues and cells, the majority of DS differentially methylated CpGs were tissue/cell-type specific. DS differentially methylated CpGs in brain tissues/cells as well as T cells were enriched for pathways related to their development and function. Another examination of DS fetal brain tissue (frontal and temporal cortex) observed a trend of modest global hypermethylation (0.3%) as well as hypermethylation of the clustered protocadherins, which are critical to neurodevelopment.^21^ Interestingly, two separate studies of DS fetal frontal cortex have found that while significant DS-differential CpGs located on HSA21 displayed a balance of hypermethylation and hypomethylation, the significant CpGs across all other chromosomes were predominantly hypermethylated.^21,22^ The DS differentially methylated CpGs belonged to genes involved in ubiquitin mediated proteolysis and Alzheimer’s disease pathways.^22^ Taken together, the differences in CpG methylation observed across a variety of DS cells/tissues are not only relevant to early developmental events but also appear to be maintained into adulthood.

Although the human DS studies discussed above have demonstrated a genome-wide signature of CpG hypermethylation in multiple DS tissues and cells, they have all made use of targeted assays that do not cover the whole genome. These targeted assays do not include regulatory elements representative of methylation across the entire genome nor do they allow for the discovery of epigenetic changes to novel regulatory regions unique to DS. Therefore, we sought to more comprehensively examine the methylomic signature of DS using WGBS and a systems biology approach to understand the impact of DNA methylation in DS cortex. Our WGBS analysis confirms and expands on the DS epigenomic literature by identifying a more comprehensive set of differentially methylated regions (DMRs) in post-mortem DS brain. Furthermore, a cross-analysis of our WGBS results with those from diverse DS tissues provides convergent evidence for reproducible genome-wide epigenetic changes associated with trisomy 21 in DS.

## Results

### WGBS analysis of DNA methylation in DS cortex reveals an abundance of hypermethylated DMRs

WGBS was performed on DNA isolated from 4 male DS and 5 male control post-mortem Broadman area 9 (BA9) samples (**Supplementary Table 1** and **Supplementary File 1**) previously described.^23^ None of the DS patients were above the age of 25 and thus are less likely to be confounded by the methylomic signature of early onset Alzheimer’s disease risk of DS. Copy number analysis of the aligned WGBS data confirmed total trisomy 21 in DS and the absence of chromosomal aneuploidies in control samples (Supplementary Figure 1). Principal component analysis (PCA) of the average methylation over 20 kb bins of CpG sites or CpG islands alone revealed no differences in large-scale DNA methylation profiles (Supplementary Figure 2). Furthermore, there were no significant differences in CpG or CpH methylation at the global level.

Given the lack of global differences in DNA methylation, we next examined locus-specific DMRs, which represent sequences spanning several hundred to a few thousand base pairs. DMRs were identified through a smoothing approach that weights CpG sites based on their coverage (Supplementary Figure 3). CpGs with correlated methylation values were grouped into DMRs, and a region statistic for each DMR was estimated, with adjustment for age (**Methods**). Statistical significance of each region was determined by permutation using a pooled null distribution. There were 9,376,534 CpG sites covered by WGBS in all samples, and 100,779 testable background regions of potential differential methylation over at least 5 contiguous CpGs (**Supplementary Table 2**). From this testable background, 3,152 significant (permutation *p* < 0.05) DMRs were identified that distinguished DS from control brain (Figure 1 and **Supplementary Table 3**). A greater number of DS-DMRs were hypermethylated (2,420) than hypomethylated (732). Interestingly, the relative abundance of hypermethylation was consistent across every chromosome (Supplementary Figure 4). A modified approach was also utilized to examine larger-scale blocks of differential methylation, where out of 87 testable blocks (**Supplementary Table 4**), 2 blocks with significant (permutation *p*<0.05) differential methylation were identified (**Supplementary Table 5** and Supplementary Figure 5). These significant blocks were not in direct proximity to protein coding genes.

**Figure 1.**
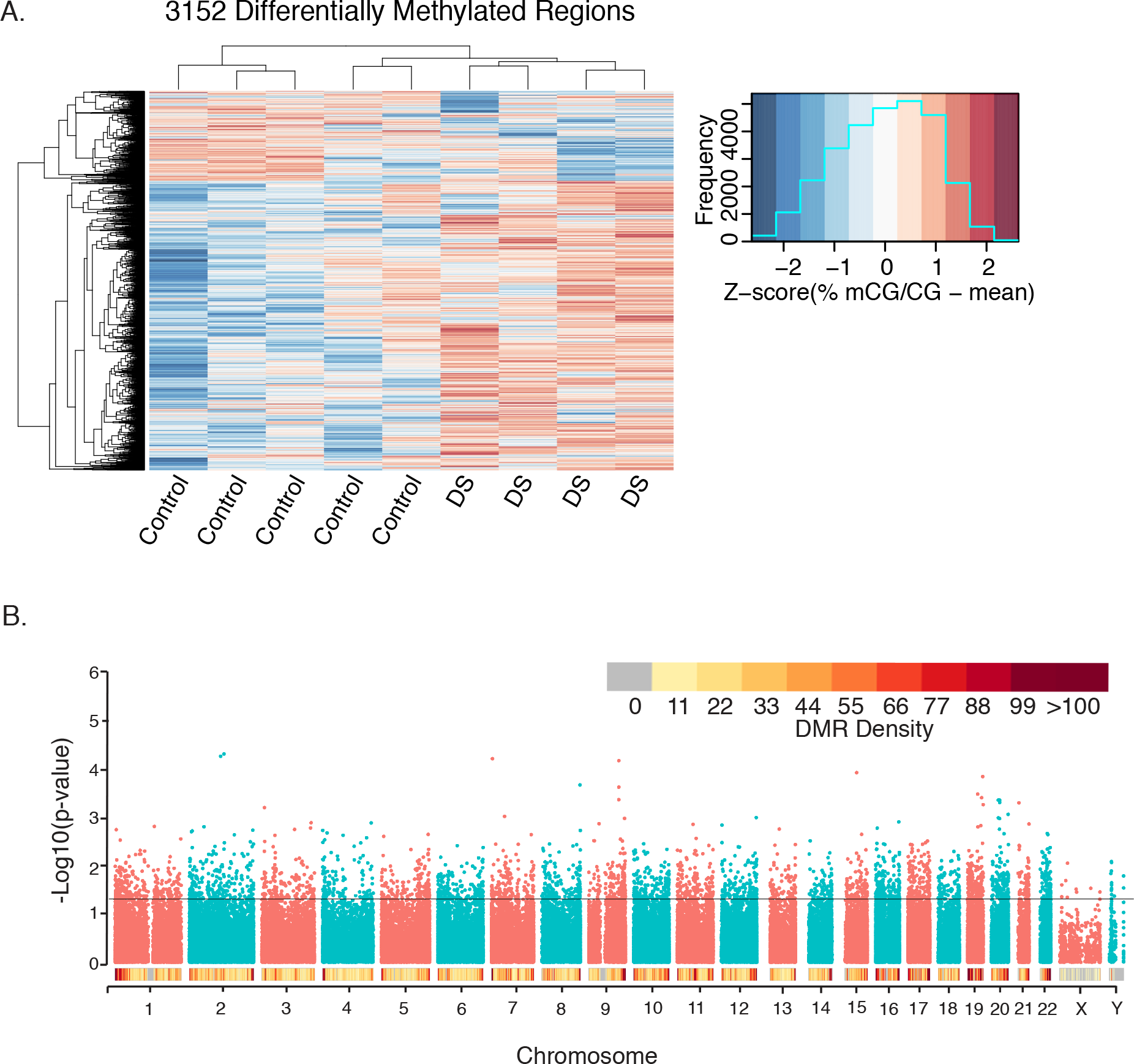
Profile of 3,152 DS-DMRs identified in post-mortem brain tissue. **A)** Heatmap visualization of hierarchical clustering analysis of DMRs. Individual values are presented as Z-scores of the smoothed percent methylation values to allow for visualization of how many standard deviations each value is from the mean. **B)** Genomic coordinate dot plot (Manhattan plot) of DS-DMRs with underlying DMR density plot of 1 Mb bins.

### Cross-tissue analyses uncover significant overlap of a subset of WGBS identified DS-DMRs

To compare the DMRs identified in our WGBS study to other genomic methylome studies in DS, we utilized previously published findings from a pan-tissue DS meta-analysis of adult brain, fetal brain, placenta, adult buccal epithelium, and adult blood samples, which identified 25 genes (24 hypermethylated and 1 hypomethylated) with differential methylation.^8^ From a gene-centric perspective, we observed a significant enrichment (Fisher’s exact test, *p*=0.0005, Odds Ratio=5.5) for 12 hypermethylated DMRs mapping to 8 genes in our WGBS identified DS-DMRs: *GLI4* (GLI Family Zinc Finger 4), *ZNF837* (Zinc Finger Protein 837), *RYR1* (Ryanodine Receptor 1), *LRRC24* (Leucine Rich Repeat Containing 24), *CELSR3* (Cadherin EGF LAG Seven-Pass G-Type Receptor), *RUNX1* (Runt Related Transcription Factor 1), *UNC45A* (Unc-45 Myosin Chaperone A), and *RFPL2* (Ret Finger Protein Like 2). Gene specific DS methylation differences are shown for six of these loci: *GLI4*, *ZNF837*, *RYR1*, *LRRC24*, *CELSR3*, and *RFPL2* (Figure 2). Notably, these pan-tissue DS-DMR associated genes were previously identified based on differential methylation of single CpG sites and are consistently identified here as larger scale DMRs.

**Figure 2.**
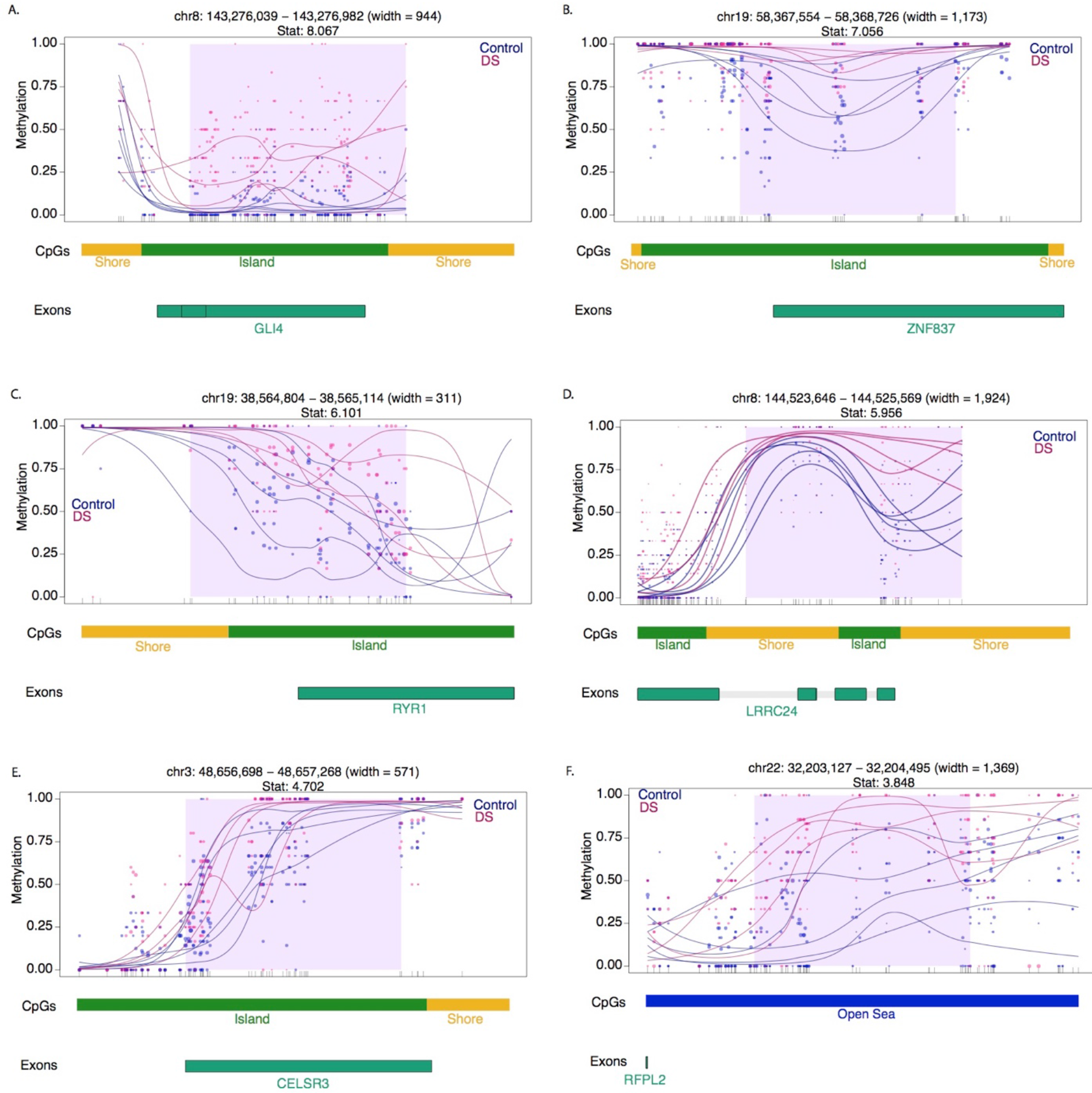
DMR plots for six select pan-tissue WGBS DS-DMRs. Each dot represents the methylation level of an individual CpG in a single sample, where the size of the dot is representative of coverage. The lines represent smoothed methylation levels for each sample, either control (blue) or DS (red). Genic and CpG annotations are shown below each plot.

In order to compare our data at the level of genomic sequence with background region and GC content accommodated for, we tested our WGBS DS-DMRs for enrichment within 12 data sets from 7 different studies that described DNA methylation differences between DS and matched control tissues (**Supplementary Table 6**). While both hypermethylated and hypomethylated categories of our WGBS DS-DMRs showed significant enrichment (*q*<0.05) with other studies across diverse tissue types, hypermethylated DS-DMRs were most frequently significantly enriched in other studies (Figure 3A). Furthermore, while there was a strong enrichment with previous studies, the majority of our WGBS-derived DS-DMRs were novel, most likely due to the increase in CpG coverage (> 9 million) of WGBS compared to the much lower CpG representation in 450k or EPIC (850k) arrays, or reduced representation bisulfite sequencing (RRBS).

**Figure 3.**
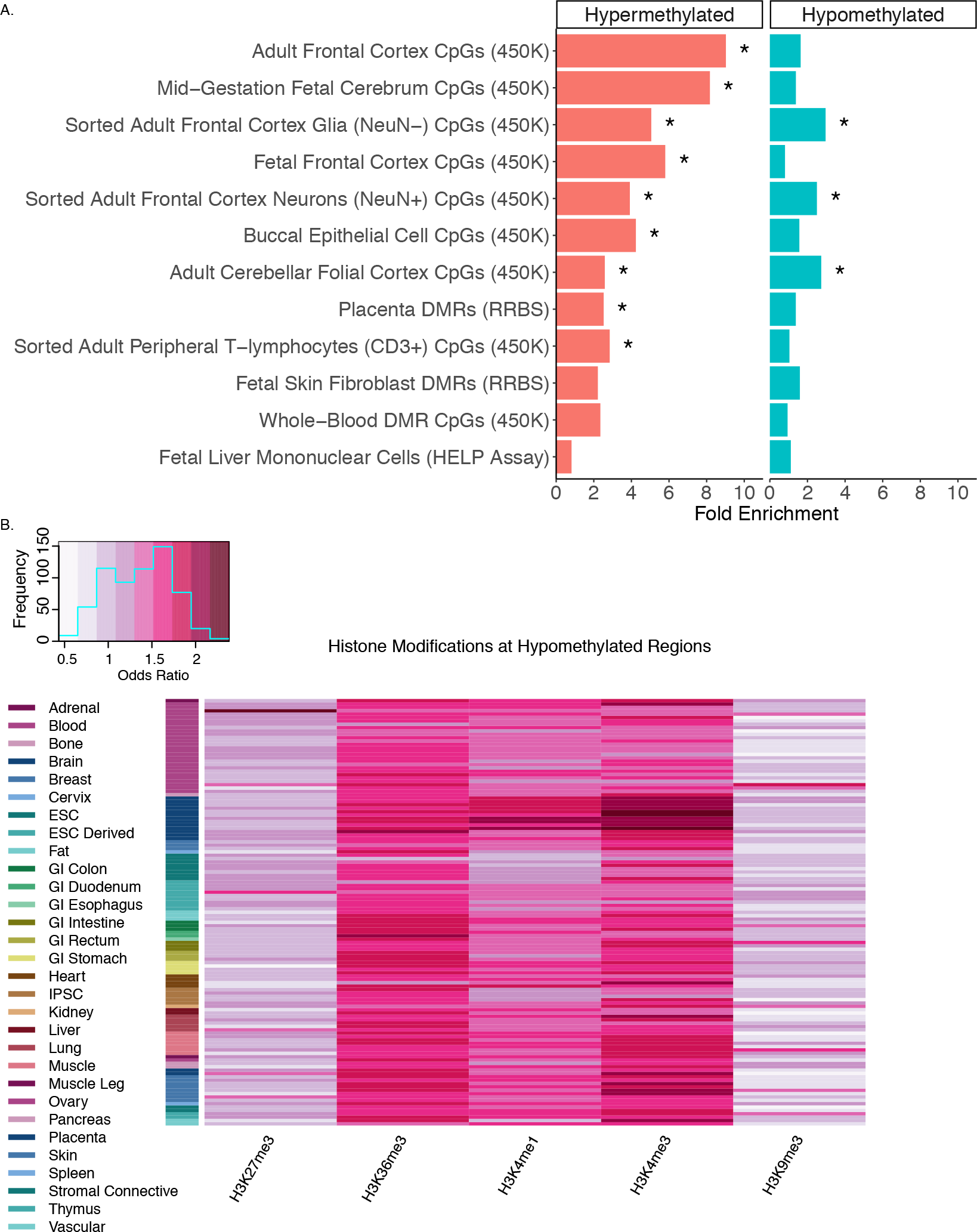
Cross-tissue enrichments for WGBS DS-DMRs relative to background regions. **A)** Overlap of DS-DMRs with differentially methylated sites and regions identified in a variety of tissues in previously published DS studies (* = *q*<0.05). **B)** Enrichment of Roadmap Epigenomics 5 core histone modifications across 127 epigenomes for the hypomethylated DS-DMRs.

Next, to further assess tissue specificity, the WGBS DS-DMRs were examined for enrichment in chromatin states (15-state model) and (5 core) histone modifications across a variety of cell types and tissues represented in the 127 Roadmap Epigenomics reference human epigenomes ^24,25^. Out of these enrichment tests (**Supplementary Table 7**), the hypomethylated DS-DMRs specifically showed significant enrichment within H3K4me3 and H3K4me1 profiles in brain tissues (Figure 3B and Supplementary Figure 6). H3K36me3 was also enriched within the hypomethylated DS-DMRs but not in a tissue-specific manner.

### Hyper- and hypo-methylated DS-DMRs affect divergent brain processes and cell types

Given the observed differences between hyper- and hypo-methylated DS-DMRs in abundance and chromatin enrichments, the DS-DMRs from WGBS were separated by directionality to perform functional annotations. When compared to background regions, both sets of DMRs were significantly (*q*<0.05) de-enriched for sequences 1-5 kb upstream of genes, promoters, introns, and intergenic regions. The hypermethylated DMRs were uniquely de-enriched for FANTOM5 permissive enhancers^26^ and 3’ UTRs of genes (Figure 4A). Furthermore, CpG based annotations revealed that both hyper- and hypomethylated DS-DMRs showed a significant (*q*<0.05) enrichment for CpG islands and a significant de-enrichment for the CpG shelves and open sea locations (Figure 4B).

**Figure 4.**
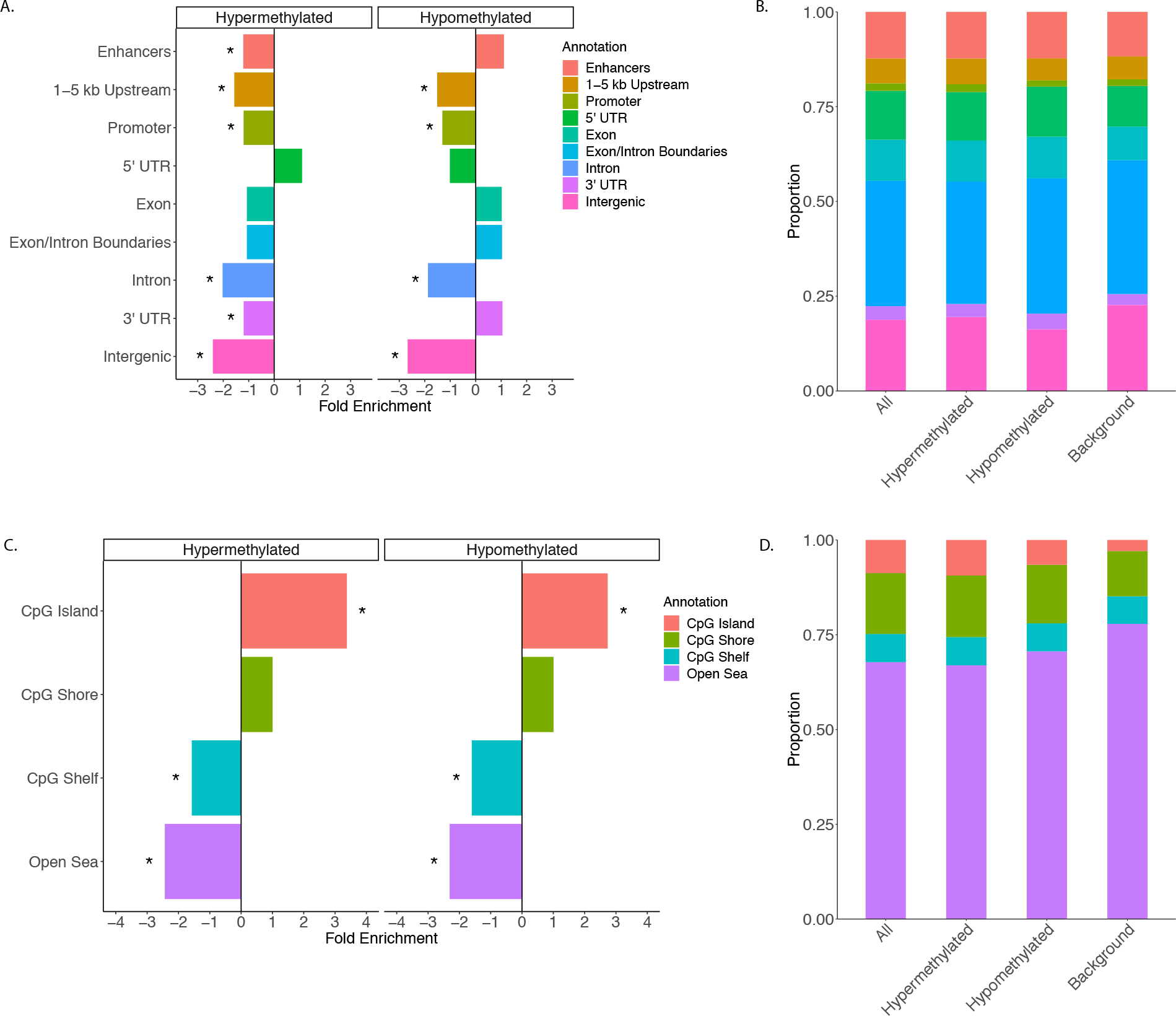
Genomic annotations of DS-DMRs relative to background regions. **A)** Enrichment of DS-DMRs in genic annotations. **B)** Proportion of DS-DMRs in genic annotations. **C)** Enrichment of DS-DMRs in CpG annotations. **D)**Proportion of DS-DMRs in CpG annotations.* = *q*<0.05 and the all category represents the combination of hypermethylated and hypomethylated DMRs.

Next, a combination of gene ontology, pathway, and transcription factor motif enrichment analyses were performed on the WGBS identified hyper- and hypomethylated DS-DMRs by utilizing genomic coordinate based approaches (Figure 5). Overall, DS-DMR associated genes were significantly (*p*<0.05) enriched for localizations to cellular membrane and its components. However, the significantly (*p*<0.05) enriched molecular and biological functions differed by directionality of DS-DMRs. The hypermethylated DS-DMRs were enriched for one-carbon metabolism, particularly through formate metabolism,^27^ and transmembrane transport functions. In contrast, hypomethylated DS-DMRs were enriched for glial immune processes and proline isomerization (Figure 5). Proline isomerization is a regulatory signaling switch with functions that span glutamatergic signaling^28^ and epigenetic modifications of the histone H3 tail that influence H3K4me3^29^ and H3K36me3^30^ levels, which are two modifications found to be significantly enriched within hypomethylated DS-DMRs (Figure 3B). The significant (*p*<0.05) canonical pathways enriched for hypermethylated DS-DMRs were primarily related to glutamatergic signaling, while hypomethylated DS-DMRs were enriched for antigen presentation and apoptosis (Figure 5). Lastly, transcription factor (TF) motif analysis revealed a significant enrichment (*p*<0.05) of 7 motifs (MYNN, PAX3:FKHR-fusion, NFAT:AP1, FRA2, E2F, CUX2, and JUN-AP1) for the hypermethylated DS-DMRs and 9 motifs (HOXA1, HOXC9, MAFK, HOXA2, GATA, DUX4, PAX7, PRDM1, and IRF1) for hypomethylated DS-DMRs (**Supplementary Table 8**). Overall, TF motifs enriched in DS-DMRs were involved in developmental processes, although the hypomethylated DS-DMRs showed additional overlap with TF motifs influencing glia. Notably, CTCF motifs were not significantly enriched within either set of DS-DMRs. Together, these results reveal multiple new insights into predicted functional consequences of trisomy 21 on epigenetic dysregulation in DS brain, extending beyond those of chromosome 21 copy number.

**Figure 5.**
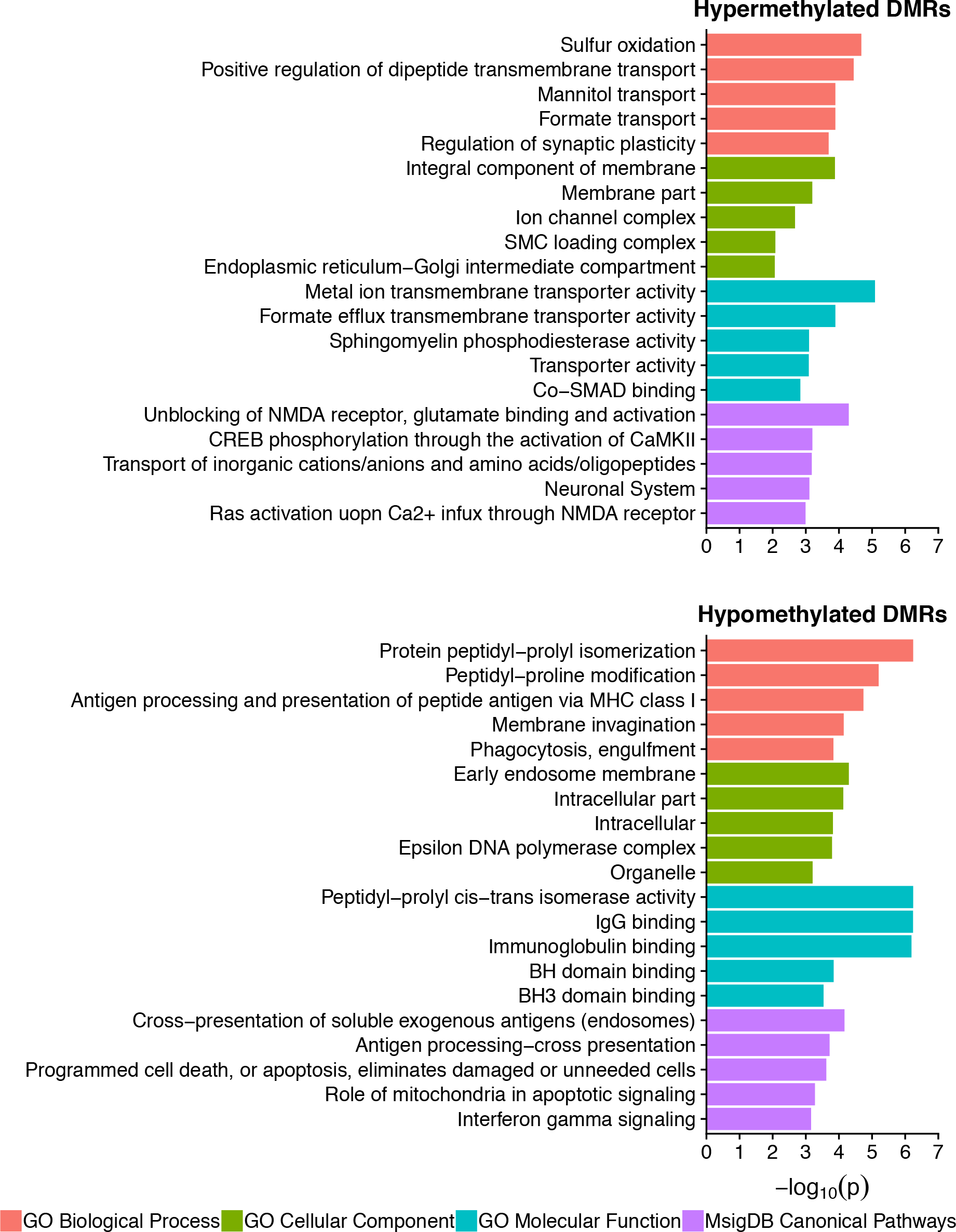
Gene ontology and pathway analysis results for hypermethylated and hypomethylated DS-DMRs relative to background regions.

## Discussion

This WGBS analysis of DMRs in DS brain confirms and extends past DS methylation studies in other tissues, where approaches using other methylation platforms showed a genome-wide impact of trisomy 21. Our results offer the most comprehensive examination of the impact of trisomy 21 on genome-wide DNA methylation patterns and identify multiple new insights into the epigenetic mechanisms of DS brain. First, the DS brain methylome was characterized by the expected abundance of hypermethylated DMRs on all chromosomes, including HSA21; however, there were no significant differences in global DNA methylation. Second, our analyses demonstrate that the hypermethylated DS-DMRs reflect both a pan-tissue signature with brain-specific impacts on glutamatergic signaling, while the hypomethylated DS-DMRs represent a more cell type specific signature related to glia and immune dysfunction. Finally, we demonstrate that both the hyper- and hypo-methylated DS-DMRs are enriched for CpG islands but de-enriched for a number of genic and regulatory regions. Overall, these results provide confirmatory evidence that trisomy 21 results in genome-wide epigenetic impacts and also offer comprehensive analyses of the impacted genes and pathways that are of relevance to DS.

Our results are consistent with several prior DS methylation analyses. Notably, a pan-tissue meta-analysis of DS tissues based on individual CpG sites identified 25 genes,^8^ 8 of which we replicated as larger scale hypermethylated DMRs in our dataset. This significant overlap was represented by 12 hypermethylated DMRs mapping to 8 different genes with functions relevant to neurodevelopment. *GLI4*, *ZNF837*, and *RUNX1* are transcription factors, while *RYR1*, *LRRC24*, *CELSR3*, *UNC45A*, and *RFPL2* are involved in signal transduction. While these 8 pan-tissue genes have been implicated in neurodevelopment through DS studies,^8,21,22^ *GLI4*,^31^ *RUNX1*,^32^ *RYR1*,^33^ *LRRC24*,^34,35^ *CELSR*,^36^ and *RFPL2*^37^ have also been shown to be involved in neurodevelopmental processes in non-DS studies, confirming the impact of DS studies on understanding general neurodevelopment. Using a more stringent genomic coordinate-based enrichment approach, we also showed strong overlaps of the hypermethylated DMRs with 12 datasets from 7 different DS methylation studies. Also consistent with the literature examining DS brain is the presence of 5 DMRs mapping to the clustered protocadherin locus,^8,20,21^ which is involved in establishing the single-cell identity of neurons.^38,39^

The relevance of our findings also extend past epigenomic studies and are complementary to those seen in a large DS plasma metabolomics study.^40^ This metabolomics study expanded on a past observation of an activated interferon response in the brain^41^ and blood^42^ of DS models, which may be explained by the presence of multiple interferon receptor genes on HSA21. Notably, interferon signaling in the brain has been shown to regulate social behavior.^43^ In the DS plasma metabolomics study, differential metabolite abundances were observed for inflammatory metabolites, L-homocysteine, the antioxidant mannitol, and sulfur metabolism.^40^ However, the most notable difference was a disrupted tryptophan catabolism that increased the levels of kynurenine and its derivative quinolinic acid, which is a neurotoxin that acts as an excitotoxic agonist of glutamatergic NMDA receptors and also plays a role in oxidative stress. The metabolites listed above are compatible with the ontologies and pathways of genes mapping to the DMRs in our study, where they are represented by differences in immune response, the one carbon metabolism (formate transport and sulfur oxidation) pathway that establishes and maintains the epigenome, mannitol transport, and glutamatergic NMDA receptors that are sensitive to the neurotoxin metabolite of the kynurenine pathway. Overall, the above similarities between the differences in metabolites and DNA methylation in DS demonstrate the utility of our approach in identifying epigenetic alterations relevant to metabolites affecting brain function. Finally, given the disruptions to similar pathways in the cortex and peripheral cells, future studies would benefit from examining neuroimmune interactions in DS.

Taken together, our WGBS results significantly extend findings from past studies by comprehensively demonstrating diverse epigenomic effects that result from a single supernumerary chromosome. Understanding how increased HSA21 copy number directly impacts methylation across the genome will require further investigation that builds on the insights gained from this and other studies. Mechanistically, there are a number of genes located on HSA21 that belong to pathways related to DNA methylation.^8^ *DNMT3L and N6AMT1* are DNA methyltransferases, while *MIS18A* interacts with DNA methyltransferases.^44^ *GART*, *CBS*, and *SOD1* are involved in one-carbon metabolism, where *SLC19A1* is a folate transporter. Functional experimentation into the cause of hypermethylation has demonstrated a key role of the DNA methyltransferase *DNMT3L*.^22^ DNMT3L is catalytically inactive; however, it is a regulatory factor that binds to and stimulates the *de novo* methyltransferases DNMT3A and DNMT3B.^45–47^ Furthermore, immunostaining suggested localization of overexpressed DNMT3L to the nuclear lamina.^22^ While these findings provide early mechanistic evidence for DNMT3L in the genome-wide profile of DNA hypermethylation, the genome-wide impact of DNMT3L overexpression remains unknown.

Our results suggest the possibility that the DS brain methylation profile arises from the hypermethylation of chromatin states accessible to additional DNMT3L molecules. Our finding of DS-specific DNA methylation in sequences enriched for H3K4me3 is supported by a previous examination of fibroblasts from DS discordant monozygotic twins showing differential gene expression occurred in late DNA replication and lamina-associated domains (LADs).^12^ While the positioning of the LADs was not altered, the active chromatin mark H3K4me3 was changed within the DS-specific chromatin domains in that study. DNMT3L is known to interact with unmethylated histone H3 to direct *de novo* methylation, which inhibits the methylation of lysine 4 (H3K4me).^48,49^ Finally, proline isomerization, which was the top observed ontology for the genes associated with hypomethylated DS-DMRs, is an epigenetic modification of the histone H3 tail that interacts with H3K4me3^29^ and H3K36me3^30^, both of which were enriched histone modifications for the hypomethylated DS-DMRs in our WGBS analysis. In order to further test the above hypotheses and address the limitations of our study, future DS WGBS studies would benefit from increased sample sizes that allow for the technical and biological validation of these results as well as the exploration of sex as a biological variable.^50^ Future DS WGBS studies would also benefit from an examination of fetal samples as well as functional examinations of *in vivo* and *in vitro* models.

Overall, our novel findings add whole-genome sequencing-level confidence to the accumulation of evidence in the literature that the brain’s entire epigenetic landscape is altered in DS in a manner that is of functional significance. Our findings extend this observation to suggest that the alterations to DNA methylation in DS reflect interactions with histone post-translational modifications. Given the dynamic nature of the epigenome, novel epigenetic editing therapeutics targeting aberrant histone or DNA methylation in DS could be attempted to improve the quality of life of patients with DS. Alternatively, the genes and pathways identified by our comprehensive approach could be useful in repurposing existing drugs for use in DS. Ultimately, the study of DS brain epigenomics is valuable not only to the DS community, but also to the broader study of other brain disorders due to the mechanistic insight into the role of DNA methylation in neurodevelopment and adult brain function.

## Materials and Methods

### Whole genome bisulfite sequencing (WGBS)

Previously published,^23^ 100 bp Illumina reads sequenced on the HiSeq 2000 of frontal cortex (Brodmann area 9; BA9) from 4 male Down Syndrome and 5 matched control samples were utilized (Sequence Read Archive: SRR3537005, SRR3537006, SRR3537007, SRR3537008, SRR3537015, SRR3537016, SRR3536978, SRR3536980, and SRX3630730). Alignments were performed using the CpG_Me pipeline (https://github.com/ben-laufer/CpG_Me). Briefly, adapters and methylation bias (m-bias) in the 5’ and 3’ ends of reads were removed from chastity filtered fastq files using Trim Galore (http://www.bioinformatics.babraham.ac.uk/projects/trim_galore/). Given that the reads were prepared by the traditional MethylC-seq library preparation protocol, a preliminary examination of methylation bias (m-bias) guided the trimming of 6 bp from the 5’ ends and 12 bp from 3’ ends. The processed reads were aligned to the human genome (hg38), deduplicated, examined for coverage, and extracted to a CpG count matrix using Bismark.^51^ Quality control and assurance (QC/QA) was performed using Trim Galore, Bismark, FastQ Screen^52^, and MultiQC.^53^

Bismark CpG count matrixes (merged cytosine reports) were then processed to generate permeth bed files of CpG methylation using the Bismark_to_Permeth_DSS.py script (https://github.com/hyeyeon-hwang/bismark-file-converter). 20kb windows and read centric CpG island windows were also created using the scripts Window_permeth_readcentric.pl and AvgMeth.2col.pl from WGBS_tools (https://github.com/kwdunaway/WGBS_Tools), respectively. Principal component analysis was performed using ggbiplot (https://github.com/vqv/ggbiplot). Global methylation was determined using a repeated measures ANOVA adjusted for age and corrected for multiple testing. Chromosome 21 copy number was determined by examining the coverage of 5 kb windows using the SAM_coverage_windowed.pl script on SAM files generated by WGBS_tools.

### Differentially methylated regions (DMRs) and blocks

DMRs were called utilizing the DMRichR workflow (https://github.com/ben-laufer/DMRichR). This workflow primarily utilizes the dmrseq^54^ and bsseq^55^ packages for inference of the DMRs and the annotatr^56^ and ChIPseeker^57^ packages for gene symbol, gene region, and CpG annotations. Briefly, CpG count matrixes (Bismark cytosine reports) were processed to merge symmetric CpG sites across strands and filtered for at least 1x coverage across samples. After filtering for coverage the majority of CpGs in the majority of samples had greater than 1x coverage (Supplementary Figure 3A), which exceeded the minimum requirements for DMR inference from WGBS data using dmrseq.^54^ DMRs were identified by testing for differences between DS BA9 brains and matched control BA9 brains, where the covariate of age was directly adjusted for. Aside from setting a cutoff of 0.05 and increasing the number of permutations to 100, DMRs were tested for using the default parameters of dmrseq. Background regions were defined as the testable regions for DMR inference, which included a methylation difference between groups, and utilized as the candidate regions for a permutation-based analysis that identified significant DMRs. Individual smoothed methylation values of the DMRs were generated using bsseq and utilized for the heatmap visualization of a hierarchal clustering analysis. Blocks of differential methylation were identified and visualized through dmrseq using the default block parameters.

### Ontology, pathway, and enrichment analyses

Genomic coordinates were lifted over to hg19 in order to perform gene ontology and pathway analysis by utilizing rGREAT (https://github.com/jokergoo/rGREAT) to access the Genomic Regions Enrichment of Annotations Tool (GREAT).^58^ For GREAT analysis, the species assembly was set to hg19 and background regions were utilized. The identified significant (unadjusted *p*<0.05) gene ontology terms were then slimed using REVIGO, where the allowed similarity was set to small (0.5) and the database with GO term sizes was set to *Homo Sapiens* (http://revigo.irb.hr).^59^

Locus Overlap Analysis (LOLA; http://databio.org/lola) was used to determine the enrichment of DMRs, relative to background regions, for reference epigenome histone modifications and chromHMM chromatin states from Core 15-state model of 5 marks 127 epigenomes (http://egg2.wustl.edu/roadmap/web_portal/chr_state_learning.html).^25^ The genomic association tester (GAT; https://github.com/AndreasHeger/gat)^60^ was used for sequence specific overlap, relative to background regions and corrected for GC content, with regulatory features and the other datasets, where the other data sets were lifted over from hg19 to hg38. 100,000 samplings were utilized for all GAT analyses. Gene symbol specific overlap was performed using GeneOverlap (https://github.com/shenlab-sinai/geneoverlap). The Hypergeometric Optimization of Motif EnRichment (HOMER) toolset (http://homer.ucsd.edu/homer/motif/) was used to identify enriched transcription factor binding sites via the findMotifsGenome.pl script, where the region size was set to size given, the normalization was set to CpG content, and the background regions were broken up to the average size of the target regions.^61^

## Acknowledgements

The authors would like to thank Keegan Korthauer from Harvard University for invaluable feedback about implementing the dmrseq analysis. The authors would also like thank Matthew Settles, Ian Korf, and Blythe Durbin-Johnson from the UC Davis Genome Center for discussions related to the bioinformatic and statistical approaches utilized in this manuscript. This work was supported by National Institutes of Health (NIH) grants [R01ES021707, R56NS076263, and U54HD079125] to J.M.L., a Canadian Institutes of Health Research (CIHR) postdoctoral fellowship [Funding Reference Number MFE-146824] to B.I.L, a NARSAD Young Investigator Award from the Brain & Behavior Research Foundation to A.V.C., and a National Institutes of Mental Health K01 Mentored Research Scientist Development award [1K01MH116389-01A1] to A.V.C.

---

**Supplementary Figure 1.**
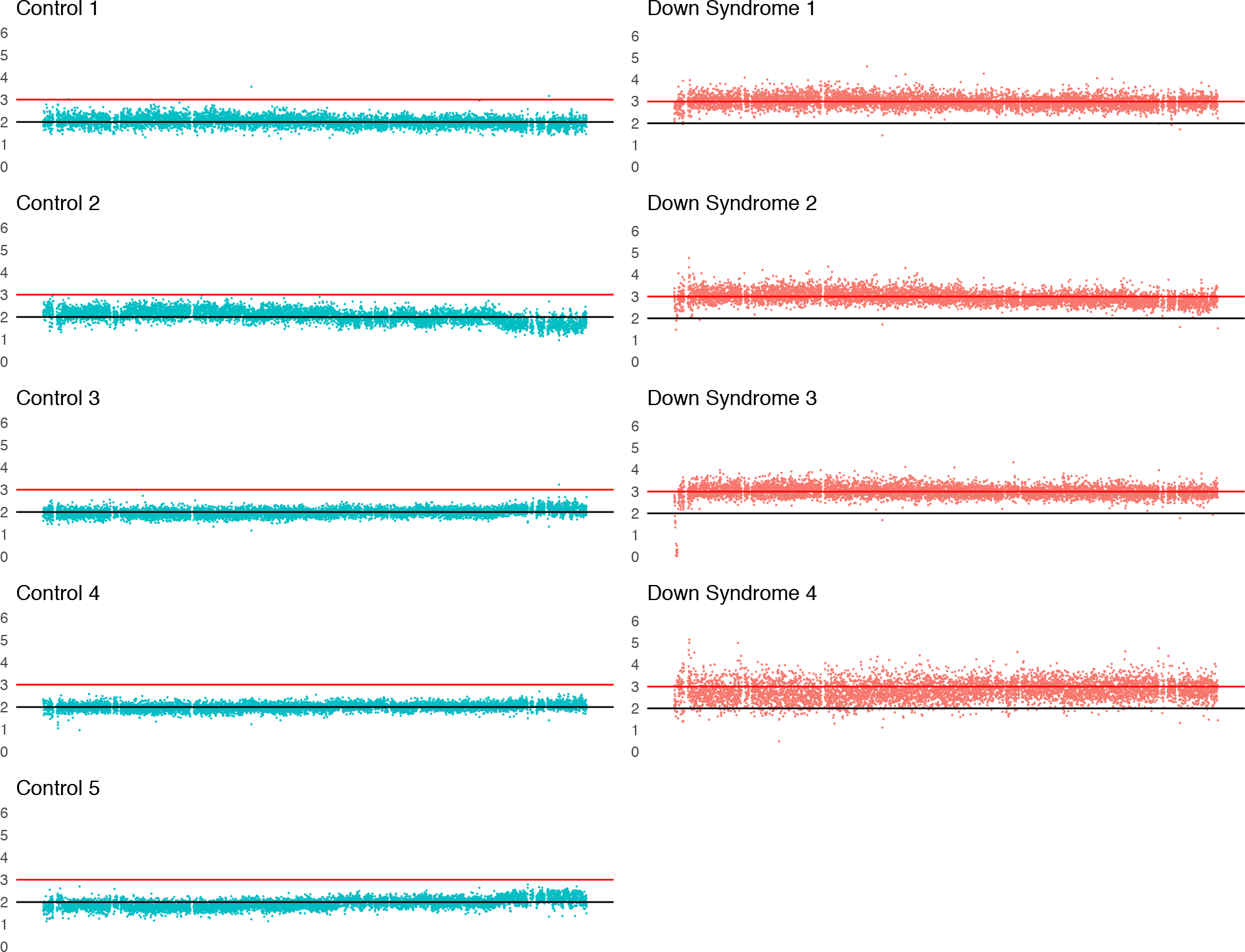
Identification of HSA21 copy number from aligned WGBS data in DS and control patients. Only the q arm of HSA21 is shown as the chromosome is acrocentric and thus reads were not aligned to the p arm due to multi-mapping.

**Supplementary Figure 2.**
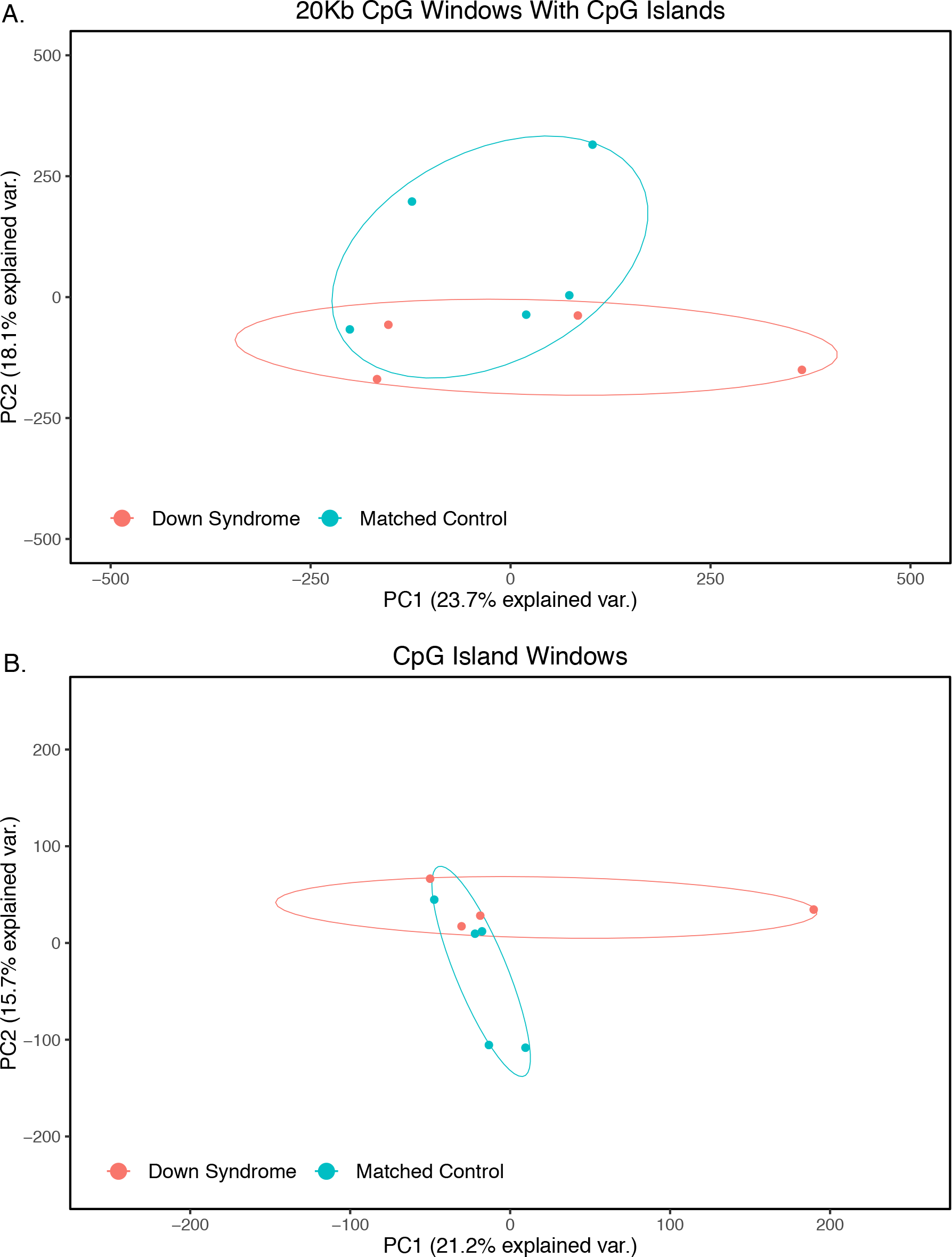
Principal component analysis of genome-wide DNA methylation in DS and control patients. **A**) 20 kb windows of average CpG methylation with CpG island sites. included. **B** Windows of average methylation across CpG islands. Ellipses represent 95% confidence intervals.

**Supplementary Figure 3.**
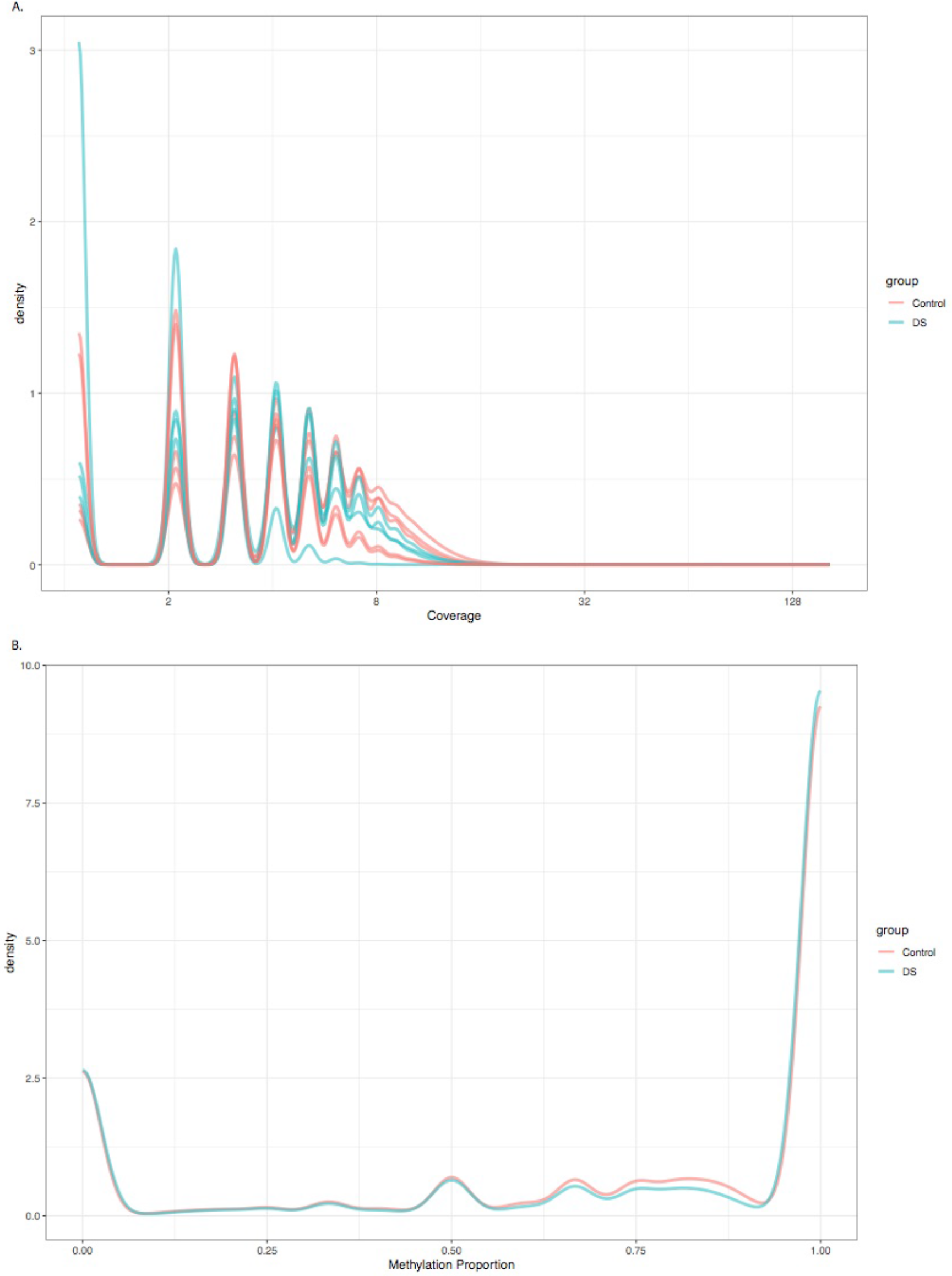
Quality control and assessment for processed CpG count matrix data used for downstream DMR analyses. **A**) Empirical distribution of CpG coverage values plotted by sample. **B**) Empirical distribution of CpG methylation values plotted by group.

**Supplementary Figure 4.**
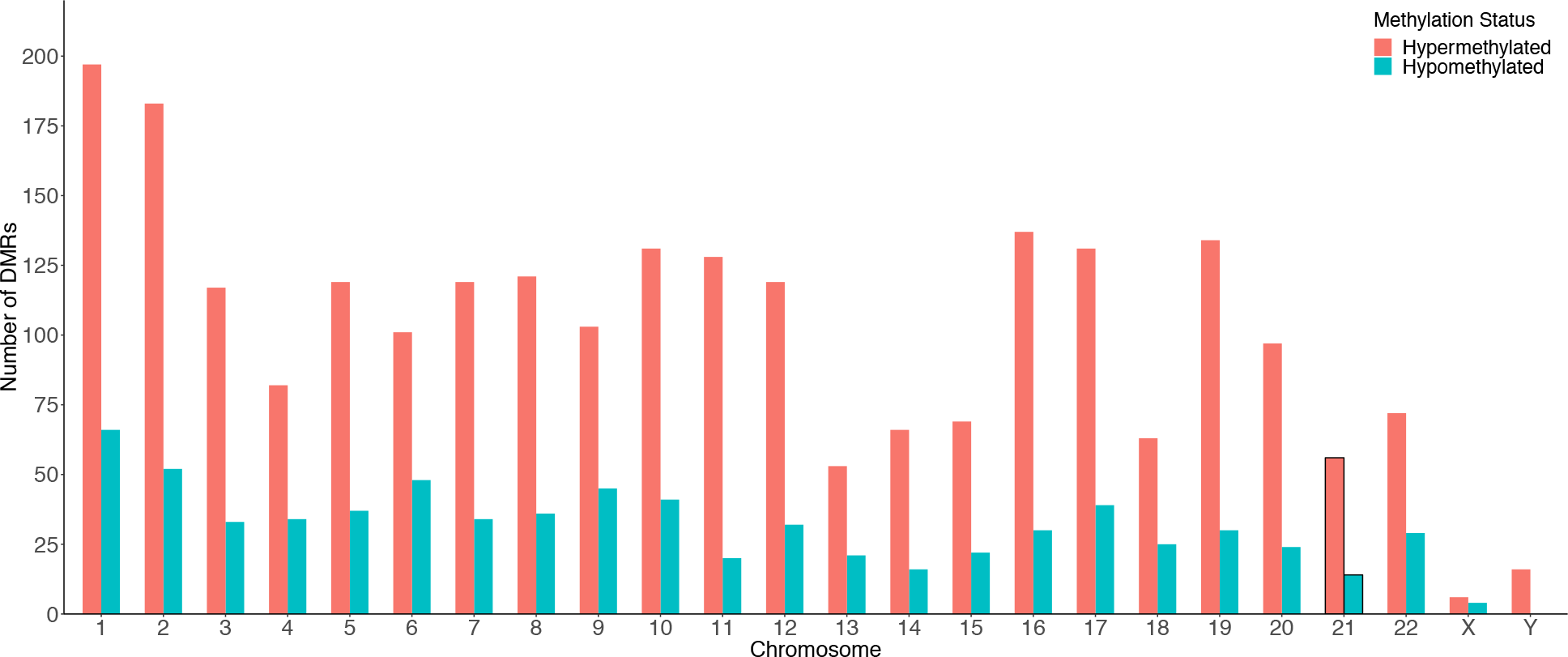
Bar plot of the distribution of DS-DMRs across the human genome.

**Supplementary Figure 5.**
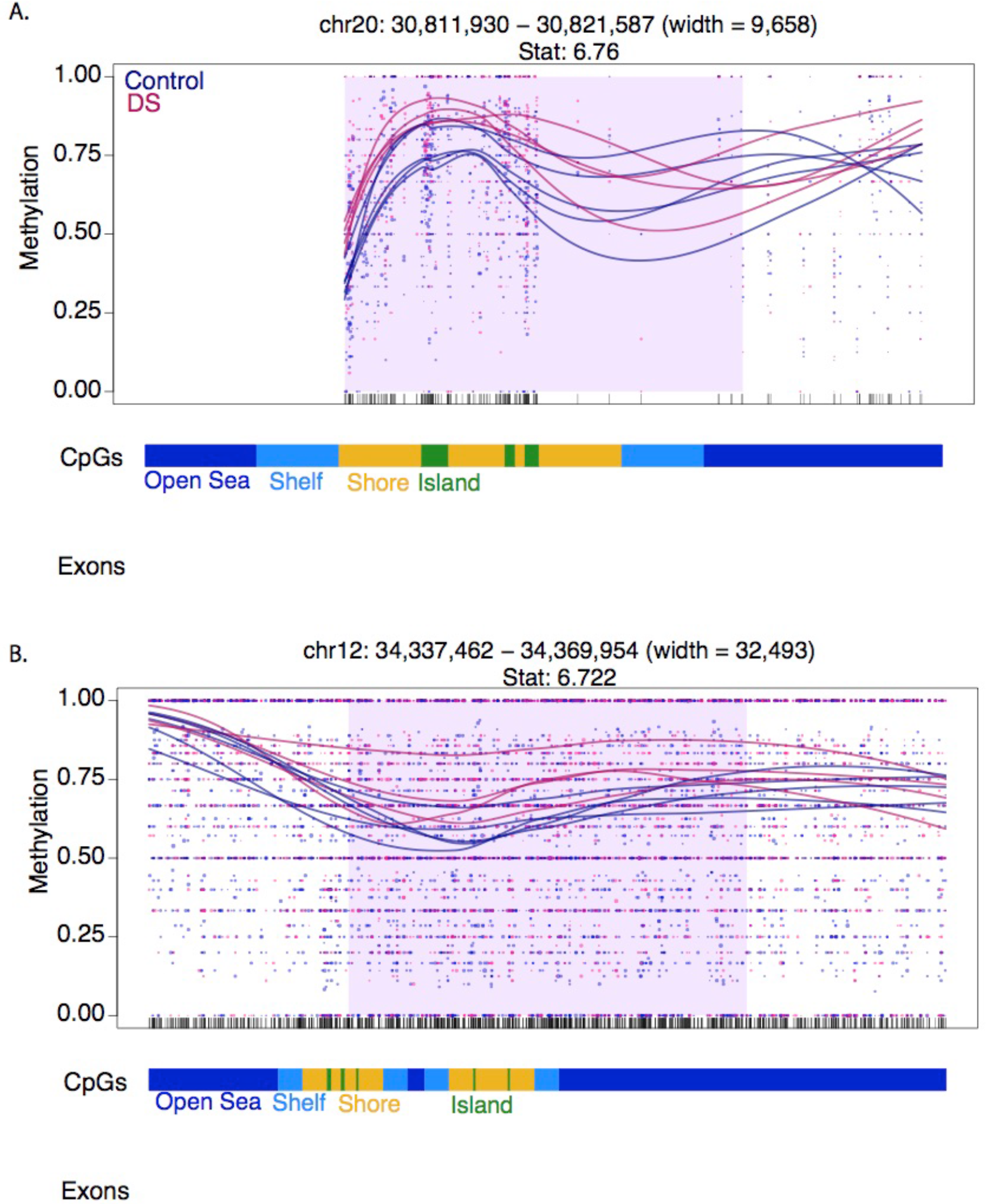
Plots of blocks of differential methylation in DS.

**Supplementary Figure 6:**
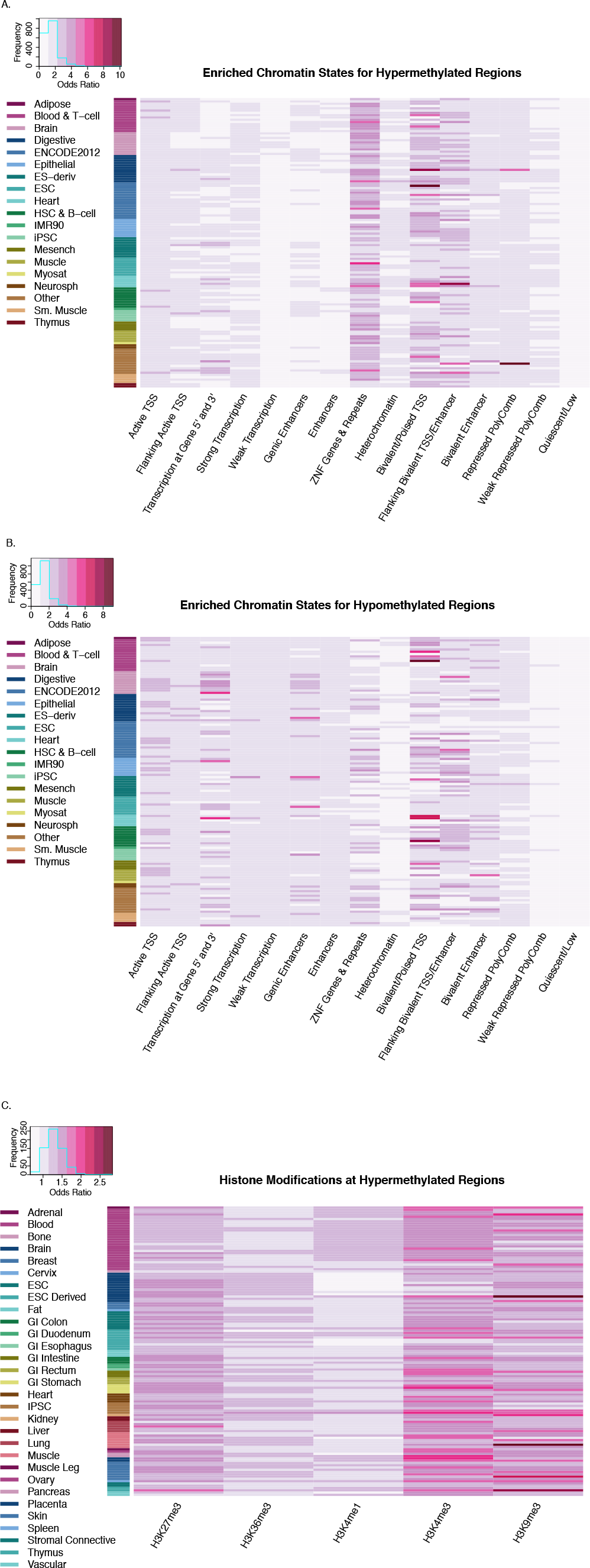
Heatmap visualization of enrichment analysis for Roadmap Epigenomics 127 epigenomes of 15-state ChromHMM model of chromatin states and 5 core histone modifications at the DS-DMRs. **A)** Chromatin states for hypermethylated DS-DMRs. **B)** Chromatin states for hypomethylated DS-DMRs. **C)** Histone modifications for hypermethylated DS-DMRs.

## Supplementary Files

Note: All Supplemental tables are provided in a single tabbed Excel spreadsheet

**Supplementary Table 1.** Post-mortem and sequencing information for DS and control samples.

**Supplementary File 1.** MultiQC report of WGBS data.

**Supplementary Table 2.** Coordinates and information for the testable background regions utilized for DMR identification.

**Supplementary Table 3.** Coordinates and information for the DMRs.

**Supplementary Figure 3.** Quality control and assessment for processed CpG count matrix data used for downstream DMR analyses. **A**) Empirical distribution of CpG coverage values plotted by sample. **B**) Empirical distribution of CpG methylation values plotted by group.

**Supplementary Table 4.** Coordinates and information for the testable background regions utilized for block identification.

**Supplementary Table 5.** Coordinates and information for blocks of differential methylation.

**Supplementary Figure 5.** Plots of blocks of differential methylation in DS.

**Supplementary Table 6:** Previously published DS data sets utilized for overlap enrichment analysis.

**Supplementary Table 7:** Enrichment analysis for Roadmap Epigenomics 127 epigenomes of 15-state ChromHMM model of chromatin states and 5 core histone modifications at the DS-DMRs. **A)** Chromatin states for hypermethylated DS-DMRs. **B)** Chromatin states for hypomethylated DS-DMRs. **C)** Histone modifications for hypermethylated DS-DMRs. **D)** Histone modifications for hypomethylated DS-DMRs.

**Supplementary Table 8:** Transcription factor motif enrichment results for known motifs within the **(A)** hypermethylated and **(B)** hypomethylated DMRs relative to background regions.

## References

1. Parker SE, Mai CT, Canfield MA, Rickard R, Wang Y, Meyer RE, Anderson P, Mason CA, Collins JS, Kirby RS, et al. Updated national birth prevalence estimates for selected birth defects in the United States, 2004-2006. Birth Defects Res Part A Clin Mol Teratol 2010; 88:1008–16.

2. Mai CT, Kucik JE, Isenburg J, Feldkamp ML, Marengo LK, Bugenske EM, Thorpe PG, Jackson JM, Correa A, Rickard R, et al. Selected birth defects data from population-based birth defects surveillance programs in the United States, 2006 to 2010: Featuring trisomy conditions. Birth Defects Res Part A Clin Mol Teratol 2013; 97:709–25.

3. Shin M, Besser LM, Kucik JE, Lu C, Siffel C, Correa A, Congenital Anomaly Multistate Prevalence and Survival Collaborative. Prevalence of Down Syndrome Among Children and Adolescents in 10 Regions of the United States. Pediatrics 2009; 124:1565–71.

4. Presson AP, Partyka G, Jensen KM, Devine OJ, Rasmussen SA, McCabe LL, McCabe ERB. Current Estimate of Down Syndrome Population Prevalence in the United States. J Pediatr 2013; 163:1163–8.

5. Korenberg JR, Chen XN, Schipper R, Sun Z, Gonsky R, Gerwehr S, Carpenter N, Daumer C, Dignan P, Disteche C. Down syndrome phenotypes: the consequences of chromosomal imbalance. Proc Natl Acad Sci U S A 1994; 91:4997–5001.

6. Yang Q, Rasmussen SA, Friedman JM. Mortality associated with Down’s syndrome in the USA from 1983 to 1997: a population-based study. Lancet (London, England) 2002; 359:1019–25.

7. Antonarakis SE. Down syndrome and the complexity of genome dosage imbalance. Nat Rev Genet 2016; 18:147–63.

8. Do C, Xing Z, Yu YE, Tycko B. Trans-acting epigenetic effects of chromosomal aneuploidies: lessons from Down syndrome and mouse models. Epigenomics 2017; 9:189–207.

9. Mao R, Zielke CL, Zielke HR, Pevsner J. Global up-regulation of chromosome 21 gene expression in the developing Down syndrome brain. Genomics 2003; 81:457–67.

10. Antonarakis SE, Lyle R, Dermitzakis ET, Reymond A, Deutsch S. Chromosome 21 and down syndrome: from genomics to pathophysiology. Nat Rev Genet 2004; 5:725–38.

11. Olmos-Serrano JL, Kang HJ, Tyler WA, Silbereis JC, Cheng F, Zhu Y, Pletikos M, Jankovic-Rapan L, Cramer NP, Galdzicki Z, et al. Down Syndrome Developmental Brain Transcriptome Reveals Defective Oligodendrocyte Differentiation and Myelination. Neuron 2016; 89:1208–22.

12. Letourneau A, Santoni FA, Bonilla X, Sailani MR, Gonzalez D, Kind J, Chevalier C, Thurman R, Sandstrom RS, Hibaoui Y, et al. Domains of genome-wide gene expression dysregulation in Down’s syndrome. Nature 2014; 508:345–50.

13. Sailani MR, Santoni FA, Letourneau A, Borel C, Makrythanasis P, Hibaoui Y, Popadin K, Bonilla X, Guipponi M, Gehrig C, et al. DNA-Methylation Patterns in Trisomy 21 Using Cells from Monozygotic Twins. PLoS One 2015; 10:e0135555.

14. Kerkel K, Schupf N, Hatta K, Pang D, Salas M, Kratz A, Minden M, Murty V, Zigman WB, Mayeux RP, et al. Altered DNA Methylation in Leukocytes with Trisomy 21. PLoS Genet 2010; 6:e1001212.

15. Bacalini MG, Gentilini D, Boattini A, Giampieri E, Pirazzini C, Giuliani C, Fontanesi E, Scurti M, Remondini D, Capri M, et al. Identification of a DNA methylation signature in blood cells from persons with Down Syndrome. Aging (Albany NY) 2014; 7:82–96.

16. Malinge S, Chlon T, Dore LC, Ketterling RP, Tallman MS, Paietta E, Gamis AS, Taub JW, Chou ST, Weiss MJ, et al. Development of acute megakaryoblastic leukemia in Down syndrome is associated with sequential epigenetic changes. Blood 2013; 122:e33–43.

17. Jones MJ, Farré P, McEwen LM, MacIsaac JL, Watt K, Neumann SM, Emberly E, Cynader MS, Virji-Babul N, Kobor MS. Distinct DNA methylation patterns of cognitive impairment and trisomy 21 in down syndrome. BMC Med Genomics 2013; 6:58.

18. Smith AK, Kilaru V, Klengel T, Mercer KB, Bradley B, Conneely KN, Ressler KJ, Binder EB. DNA extracted from saliva for methylation studies of psychiatric traits: Evidence tissue specificity and relatedness to brain. Am J Med Genet Part B Neuropsychiatr Genet 2015; 168:36–44.

19. Jin S, Lee YK, Lim YC, Zheng Z, Lin XM, Ng DPY, Holbrook JD, Law HY, Kwek KYC, Yeo GSH, et al. Global DNA Hypermethylation in Down Syndrome Placenta. PLoS Genet 2013; 9:e1003515.

20. Mendioroz M, Do C, Jiang X, Liu C, Darbary HK, Lang CF, Lin J, Thomas A, Abu-Amero S, Stanier P, et al. Trans effects of chromosome aneuploidies on DNA methylation patterns in human Down syndrome and mouse models. Genome Biol 2015; 16:263.

21. El Hajj N, Dittrich M, Böck J, Kraus TFJ, Nanda I, Müller T, Seidmann L, Tralau T, Galetzka D, Schneider E, et al. Epigenetic dysregulation in the developing Down syndrome cortex. Epigenetics 2016; 11:563–78.

22. Lu J, Mccarter M, Lian G, Esposito G, Capoccia E, Delli-Bovi LC, Hecht J, Sheen V. Global hypermethylation in fetal cortex of Down syndrome due to DNMT3L overexpression. Hum Mol Genet 2016; 25:1714–27.

23. Dunaway KW, Islam MS, Coulson RL, Lopez SJ, Vogel Ciernia A, Chu RG, Yasui DH, Pessah IN, Lott P, Mordaunt C, et al. Cumulative Impact of Polychlorinated Biphenyl and Large Chromosomal Duplications on DNA Methylation, Chromatin, and Expression of Autism Candidate Genes. Cell Rep 2016; 17:3035–48.

24. Ernst J, Kellis M. ChromHMM: automating chromatin-state discovery and characterization. Nat Methods 2012; 9:215–6.

25. Roadmap Epigenomics Consortium, Kundaje A, Meuleman W, Ernst J, Bilenky M, Yen A, Heravi-Moussavi A, Kheradpour P, Zhang Z, Wang J, et al. Integrative analysis of 111 reference human epigenomes. Nature 2015; 518:317–30.

26. Andersson R, Gebhard C, Miguel-Escalada I, Hoof I, Bornholdt J, Boyd M, Chen Y, Zhao X, Schmidl C, Suzuki T, et al. An atlas of active enhancers across human cell types and tissues. Nature 2014; 507:455–61.

27. Brosnan ME, Brosnan JT. Formate: The Neglected Member of One-Carbon Metabolism. Annu Rev Nutr 2016; 36:369–88.

28. Antonelli R, De Filippo R, Middei S, Stancheva S, Pastore B, Ammassari-Teule M, Barberis A, Cherubini E, Zacchi P. Pin1 Modulates the Synaptic Content of NMDA Receptors via Prolyl-Isomerization of PSD-95. J Neurosci 2016; 36:5437–47.

29. Howe FS, Boubriak I, Sale MJ, Nair A, Clynes D, Grijzenhout A, Murray SC, Woloszczuk R, Mellor J. Lysine Acetylation Controls Local Protein Conformation by Influencing Proline Isomerization. Mol Cell 2014; 55:733–44.

30. Nelson CJ, Santos-Rosa H, Kouzarides T. Proline isomerization of histone H3 regulates lysine methylation and gene expression. Cell 2006; 126:905–16.

31. Laurendeau I, Ferrer M, Garrido D, D’Haene N, Ciavarelli P, Basso A, Vidaud M, Bieche I, Salmon I, Szijan I. Gene expression profiling of the hedgehog signaling pathway in human meningiomas. Mol Med 2010; 16:262–70.

32. Inoue K, Shiga T, Ito Y. Runx transcription factors in neuronal development. Neural Dev 2008; 3:20.

33. Wayman GA, Yang D, Bose DD, Lesiak A, Ledoux V, Bruun D, Pessah IN, Lein PJ. PCB-95 promotes dendritic growth via ryanodine receptor-dependent mechanisms. Environ Health Perspect 2012; 120:997–1002.

34. Söllner C, Wright GJ. A cell surface interaction network of neural leucine-rich repeat receptors. Genome Biol 2009; 10:R99.

35. Gilsohn E, Volk T. Fine tuning cellular recognition: The function of the leucine rich repeat (LRR) trans-membrane protein, LRT, in muscle targeting to tendon cells. Cell Adh Migr 2010; 4:368–71.

36. Tissir F, Bar I, Jossin Y, De Backer O, Goffinet AM. Protocadherin Celsr3 is crucial in axonal tract development. Nat Neurosci 2005; 8:451–7.

37. Bonnefont J, Nikolaev SI, Perrier AL, Guo S, Cartier L, Sorce S, Laforge T, Aubry L, Khaitovich P, Peschanski M, et al. Evolutionary Forces Shape the Human RFPL1,2,3 Genes toward a Role in Neocortex Development. Am J Hum Genet 2008; 83:208–18.

38. Chen W V, Maniatis T. Clustered protocadherins. Development 2013; 140:3297–302.

39. Hayashi S, Takeichi M. Emerging roles of protocadherins: from self-avoidance to enhancement of motility. J Cell Sci 2015; 128:1455–64.

40. Powers RK, Sullivan KD, Culp-Hill R, Ludwig MP, Smith KP, Waugh KA, Minter R, Tuttle KD, Rachubinski AL, Granrath RE, et al. Trisomy 21 drives production of neurotoxic tryptophan catabolites via the interferon-inducible kynurenine pathway. bioRxiv 2018; :403642.

41. Aziz NM, Guedj F, Pennings JLA, Olmos-Serrano JL, Siegel A, Haydar TF, Bianchi DW. Lifespan analysis of brain development, gene expression and behavioral phenotypes in the Ts1Cje, Ts65Dn and Dp(16)1/Yey mouse models of Down syndrome. Dis Model Mech 2018; 11:dmm031013.

42. Sullivan KD, Lewis HC, Hill AA, Pandey A, Jackson LP, Cabral JM, Smith KP, Liggett LA, Gomez EB, Galbraith MD, et al. Trisomy 21 consistently activates the interferon response. Elife 2016; 5.

43. Filiano AJ, Xu Y, Tustison NJ, Marsh RL, Baker W, Smirnov I, Overall CC, Gadani SP, Turner SD, Weng Z, et al. Unexpected role of interferon-γ in regulating neuronal connectivity and social behaviour. Nature 2016; 535:425–9.

44. Kim IS, Lee M, Park KC, Jeon Y, Park JH, Hwang EJ, Jeon TI, Ko S, Lee H, Baek SH, et al. Roles of Mis18α in Epigenetic Regulation of Centromeric Chromatin and CENP-A Loading. Mol Cell 2012; 46:260–73.

45. Chedin F, Lieber MR, Hsieh C-L. The DNA methyltransferase-like protein DNMT3L stimulates de novo methylation by Dnmt3a. Proc Natl Acad Sci 2002; 99:16916–21.

46. Chen Z-X, Mann JR, Hsieh C-L, Riggs AD, Chédin F. Physical and functional interactions between the human DNMT3L protein and members of the de novo methyltransferase family. J Cell Biochem 2005; 95:902–17.

47. Jeltsch A, Jurkowska RZ. Allosteric control of mammalian DNA methyltransferases – a new regulatory paradigm. Nucleic Acids Res 2016; 44:8556–75.

48. Ooi SKT, Qiu C, Bernstein E, Li K, Jia D, Yang Z, Erdjument-Bromage H, Tempst P, Lin S-P, Allis CD, et al. DNMT3L connects unmethylated lysine 4 of histone H3 to de novo methylation of DNA. Nature 2007; 448:714–7.

49. Otani J, Nankumo T, Arita K, Inamoto S, Ariyoshi M, Shirakawa M. Structural basis for recognition of H3K4 methylation status by the DNA methyltransferase 3A ATRX-DNMT3-DNMT3L domain. EMBO Rep 2009; 10:1235–41.

50. Ratnu VS, Emami MR, Bredy TW. Genetic and epigenetic factors underlying sex differences in the regulation of gene expression in the brain. J Neurosci Res 2017; 95:301–10.

51. Krueger F, Andrews SR. Bismark: a flexible aligner and methylation caller for Bisulfite-Seq applications. Bioinformatics 2011; 27:1571–2.

52. Wingett SW, Andrews S. FastQ Screen: A tool for multi-genome mapping and quality control. F1000Research 2018; 7:1338.

53. Ewels P, Magnusson M, Lundin S, Käller M. MultiQC: summarize analysis results for multiple tools and samples in a single report. Bioinformatics 2016; 32:3047–8.

54. Korthauer K, Chakraborty S, Benjamini Y, Irizarry RA. Detection and accurate false discovery rate control of differentially methylated regions from whole genome bisulfite sequencing. Biostatistics 2018;

55. Hansen KD, Langmead B, Irizarry RA. BSmooth: from whole genome bisulfite sequencing reads to differentially methylated regions. Genome Biol 2012; 13:R83.

56. Cavalcante RG, Sartor MA. annotatr: genomic regions in context. Bioinformatics 2017; 33:2381–3.

57. Yu G, Wang L-G, He Q-Y. ChIPseeker: an R/Bioconductor package for ChIP peak annotation, comparison and visualization. Bioinformatics 2015; 31:2382–3.

58. McLean CY, Bristor D, Hiller M, Clarke SL, Schaar BT, Lowe CB, Wenger AM, Bejerano G. GREAT improves functional interpretation of cis-regulatory regions. Nat Biotechnol 2010; 28:495–501.

59. Supek F, Bošnjak M, Škunca N, Šmuc T. REVIGO summarizes and visualizes long lists of gene ontology terms. PLoS One 2011; 6:e21800.

60. Heger A, Webber C, Goodson M, Ponting CP, Lunter G. GAT: a simulation framework for testing the association of genomic intervals. Bioinformatics 2013; 29:2046–8.

61. Heinz S, Benner C, Spann N, Bertolino E, Lin YC, Laslo P, Cheng JX, Murre C, Singh H, Glass CK. Simple Combinations of Lineage-Determining Transcription Factors Prime cis-Regulatory Elements Required for Macrophage and B Cell Identities. Mol Cell 2010; 38:576–89

