## Supplementary File 1 for "Whole genome bisulfite sequencing of Down syndrome brain reveals regional DNA hypermethylation and novel disorder insights"

CpG\_Me: MultiQC Report


### Toggle navigation v1.5

### CpG\_Me

Loading report..

- General Stats
- Bismark
  - Alignment Rates
  - Deduplication
  - Strand Alignment
  - Cytosine Methylation
  - M-Bias
- FastQ Screen
- FastQC
  - Sequence Quality Histograms
  - Per Sequence Quality Scores
  - Per Base Sequence Content
  - Per Sequence GC Content
  - Per Base N Content
  - Sequence Length Distribution
  - Sequence Duplication Levels
  - Overrepresented sequences
  - Adapter Content
- Cutadapt

Toolbox

##### MultiQC Toolbox

###### Apply Highlight Samples

+

Regex mode off
help
 Clear

###### Apply Rename Samples

+

Click here for bulk input.

Paste two columns of a tab-delimited table here (eg. from Excel).

First column should be the old name, second column the new name.

Add

Regex mode off
help
 Clear

###### Apply Show / Hide Samples

Hide matching samples

Show only matching samples

+

Regex mode off
help
 Clear

###### Export Plots

- Images
- Data

px

px

Aspect ratio

PNG
JPEG
SVG

Plot scaling

X

Download the raw data used to create the plots in this report below:

Format:

Tab-separated
Comma-separated
JSON

Note that additional data was saved in `multiqc_data` when this report was generated.

---

###### Choose Plots

 All
 None


---


   Download Plot Images

If you use plots from MultiQC in a publication or presentation, please cite:

> **MultiQC: Summarize analysis results for multiple tools and samples in a single report**  
> *Philip Ewels, Måns Magnusson, Sverker Lundin and Max Käller*  
> Bioinformatics (2016)  
> doi: 10.1093/bioinformatics/btw354  
> PMID: 27312411

###### Save Settings

You can save the toolbox settings for this report to the browser.

 Save


---

###### Load Settings

Choose a saved report profile from the dropdown box below:

[ select ]

Load
 Delete
 Set default
 Clear default

###### About MultiQC

This report was generated using MultiQC, version 1.5

You can see a YouTube video describing how to use MultiQC reports here:
https://youtu.be/qPbIlO\_KWN0

For more information about MultiQC, including other videos and
extensive documentation, please visit http://multiqc.info

You can report bugs, suggest improvements and find the source code for MultiQC on GitHub:
https://github.com/ewels/MultiQC

MultiQC is published in Bioinformatics:

> **MultiQC: Summarize analysis results for multiple tools and samples in a single report**  
> *Philip Ewels, Måns Magnusson, Sverker Lundin and Max Käller*  
> Bioinformatics (2016)  
> doi: 10.1093/bioinformatics/btw354  
> PMID: 27312411

### CpG\_Me QC for a custom WGBS workflow

Multiple QC reports summarise filtering, trimming, alignment, and methylation bias (m-bias) results.

> This is the LaSalle lab version of CpG\_Me for single end sequencing.

Workflow Developer
:   Ben Laufer

E-mail
:  

Application Type
:   WGBS

Project Type
:   Down Syndrome

Sequencing Platform
:   HiSeq 2000

Sequencing Setup
:   SE 100 SI

Library Kit
:   MethylC-seq

Genome
:   hg38

###### JavaScript Disabled

MultiQC reports use JavaScript for plots and toolbox functions. It looks like
you have JavaScript disabled in your web browser. Please note that many of the report
functions will not work as intended.

Loading report..

Report generated on 2018-06-03, 11:03 based on data in:
`/share/lasallelab/Ben/DownSyndrome/Update`

---

×
don't show again

**Welcome!** Not sure where to start?  
Watch a tutorial video
  *(6:06)*

#### General Statistics

 Copy table

 Configure Columns

 Sort by highlight

 Plot
Showing ^9^/_9_ rows and ^8^/_13_ columns.

| Sample Name | % mCpG | M C's | C Coverage | % Dups | M Unique | M Aligned | % Aligned | % Dups | % GC | Length | % Failed | M Seqs | % Trimmed |
| --- | --- | --- | --- | --- | --- | --- | --- | --- | --- | --- | --- | --- | --- |
| JLKD062\_filtered | 76.2% | 1630.4 | 2.60X | 6.4% | 119.5 | 127.7 | 71.1% | 9.7% | 28% | 68 bp | 9% | 179.6 | 15.5% |
| SRR3536978 | 76.8% | 1566.2 | 2.50X | 6.0% | 114.9 | 122.2 | 74.3% | 11.5% | 27% | 67 bp | 9% | 164.4 | 1.7% |
| SRR3536980 | 77.2% | 1561.4 | 2.49X | 4.1% | 118.1 | 123.2 | 72.0% | 10.1% | 26% | 66 bp | 9% | 171.2 | 1.6% |
| SRR3537005 | 74.0% | 1181.2 | 1.89X | 4.6% | 99.0 | 103.7 | 71.8% | 12.7% | 27% | 62 bp | 9% | 144.5 | 2.3% |
| SRR3537006 | 78.4% | 1502.1 | 2.40X | 5.4% | 108.5 | 114.7 | 75.2% | 10.6% | 26% | 69 bp | 18% | 152.5 | 1.5% |
| SRR3537007 | 78.1% | 1540.2 | 2.46X | 4.1% | 121.3 | 126.5 | 72.9% | 9.8% | 26% | 63 bp | 9% | 173.6 | 2.0% |
| SRR3537008 | 74.8% | 434.7 | 0.69X | 17.8% | 28.0 | 34.1 | 77.3% | 21.1% | 27% | 78 bp | 18% | 44.0 | 2.1% |
| SRR3537015 | 74.8% | 1014.4 | 1.61X | 4.8% | 82.3 | 86.4 | 56.6% | 9.6% | 25% | 65 bp | 9% | 152.6 | 2.0% |
| SRR3537016 | 76.8% | 1020.9 | 1.62X | 3.5% | 81.2 | 84.1 | 59.0% | 8.8% | 25% | 66 bp | 9% | 142.6 | 1.7% |

×

###### General Statistics: Columns

Uncheck the tick box to hide columns. Click and drag the handle on the left to change order.

Show All
Show None

| Sort | Visible | Group | Column | Description | ID | Scale |
| --- | --- | --- | --- | --- | --- | --- |
| \|\| |  | Bismark | % mCpG | % Cytosines methylated in CpG context | `percent_cpg_meth` | None |
| \|\| |  | Bismark | M C's | Total number of C's analysed, in millions | `total_c` | None |
| \|\| |  | Bismark | C Coverage | Cyotosine Coverage | `C_coverage` | None |
| \|\| |  | Bismark | % Dups | Percent Duplicated Alignments | `dup_reads_percent` | None |
| \|\| |  | Bismark | M Unique | Deduplicated Alignments (millions) | `dedup_reads` | read\_count |
| \|\| |  | Bismark | M Aligned | Total Aligned Sequences (millions) | `aligned_reads` | read\_count |
| \|\| |  | Bismark | % Aligned | Percent Aligned Sequences | `percent_aligned` | None |
| \|\| |  | FastQC | % Dups | % Duplicate Reads | `percent_duplicates` | None |
| \|\| |  | FastQC | % GC | Average % GC Content | `percent_gc` | None |
| \|\| |  | FastQC | Length | Average Sequence Length (bp) | `avg_sequence_length` | None |
| \|\| |  | FastQC | % Failed | Percentage of modules failed in FastQC report (includes those not plotted here) | `percent_fails` | None |
| \|\| |  | FastQC | M Seqs | Total Sequences (millions) | `total_sequences` | read\_count |
| \|\| |  | Cutadapt | % Trimmed | % Total Base Pairs trimmed | `percent_trimmed` | None |

Close

#### Bismark

Bismark is a tool to map bisulfite converted sequence reads and determine cytosine methylation states.

##### Alignment Rates

Number of Reads
Percentages

loading..

---

##### Deduplication

Number of Reads
Percentages

loading..

---

##### Strand Alignment

All samples were run with `--directional` mode; alignments to complementary strands (CTOT, CTOB) were ignored.

Number of Reads
Percentages

loading..

---

##### Cytosine Methylation

loading..

---

##### M-Bias

This plot shows the average percentage methylation and coverage across reads. See the
bismark user guide
for more information on how these numbers are generated.

CpG R1
CHG R1
CHH R1

loading..

---

#### FastQ Screen

FastQ Screen allows you to screen a library of sequences in FastQ format against a set of sequence databases so you can see if the composition of the library matches with what you expect.

---

#### FastQC

FastQC is a quality control tool for high throughput sequence data, written by Simon Andrews at the Babraham Institute in Cambridge.

##### Sequence Quality Histograms

The mean quality value across each base position in the read. See the FastQC help.

loading..

---

##### Per Sequence Quality Scores

The number of reads with average quality scores. Shows if a subset of reads has poor quality. See the FastQC help.

loading..

---

##### Per Base Sequence Content

The proportion of each base position for which each of the four normal DNA bases has been called. See the FastQC help.

Click a sample row to see a line plot for that dataset.

###### Rollover for sample name

 Export Plot

Position: -

%T: -

%C: -

%A: -

%G: -

---

##### Per Sequence GC Content

The average GC content of reads. Normal random library typically have a roughly normal distribution of GC content.
See the FastQC help.

Percentages
Counts

loading..

---

##### Per Base N Content

The percentage of base calls at each position for which an N was called. See the FastQC help.

loading..

---

##### Sequence Length Distribution

The distribution of fragment sizes (read lengths) found. See the FastQC help.

loading..

---

##### Sequence Duplication Levels

The relative level of duplication found for every sequence. See the FastQC help.

loading..

---

##### Overrepresented sequences

The total amount of overrepresented sequences found in each library. See the FastQC help for further information.

9 samples had less than 1% of reads made up of overrepresented sequences

---

##### Adapter Content

The cumulative percentage count of the proportion of your library which has seen each of the adapter sequences at each position. See the FastQC help. Only samples with ≥ 0.1% adapter contamination are shown.

No samples found with any adapter contamination > 0.1%

---

#### Cutadapt

Cutadapt is a tool to find and remove adapter sequences, primers, poly-Atails and other types of unwanted sequence from your high-throughput sequencing reads.

This plot shows the number of reads with certain lengths of adapter trimmed.
Obs/Exp shows the raw counts divided by the number expected due to sequencing errors. A defined peak
may be related to adapter length. See the
cutadapt documentation
for more information on how these numbers are generated.

Counts
Obs/Exp

loading..

**MultiQC v1.5** 
- Written by Phil Ewels,
available on GitHub.

This report uses HighCharts,
jQuery,
jQuery UI,
Bootstrap,
FileSaver.js and
clipboard.js.

×

##### Plot Table Data

Select Column

Select Column

Please select two table columns.

Close

×

##### Regex Help

Toolbox search strings can behave as regular expressions (regexes). Click a button below to see an example of it in action. Try modifying them yourself in the text box.

`^` (start of string)
`$` (end of string)
`[]` (character choice)
`\d` (shorthand for `[0-9]`)
`\w` (shorthand for `[0-9a-zA-Z_]`)
`.` (any character)
`\.` (literal full stop)
`()` `|` (group / separator)
`*` (prev char 0 or more)
`+` (prev char 1 or more)
`?` (prev char 0 or 1)
`{}` (char num times)
`{,}` (count range)

```
samp_1
samp_1_edited
samp_2
samp_2_edited
samp_3
samp_3_edited
prepended_samp_1
tmp_samp_1_edited
tmpp_samp_1_edited
tmppp_samp_1_edited
#samp_1_edited.tmp
samp_11
samp_11111
```

See regex101.com for a more heavy duty testing suite.

Close
